# HEPG2 EXPRESSION OF MIR-202 & TRIB-1 UNDER METABOLIC AND INFLAMMATORY STRESS

**DOI:** 10.1101/2021.07.29.454303

**Authors:** Iquo O. Phillip, Julius O. Phillip

## Abstract

**Background:** The burden of cardiovascular disease (CVD) is such that affects both developed and developing countries with high rates of mortality and morbidity. Cardiovascular diseases are highly polymorphic across its various risk factors. Human polymorphisms of trib-1 gene have been implicated to be associated with risk factors for CVD. Trib-1 gene is a known target for microRNA-202 which consequently could have an effect on its stability. The objective of this study was to evaluate the expression of miR-202 in an hepatic cell line under *in vitro* conditions of metabolic and inflammatory stress and the effect on trib-1 level.

**Materials and Methods:** HepG2 cells cultured under *in vitro* conditions of high glucose and cytokine stimulation of concentrations of varying time intervals were harvested and mRNA/microRNA extracted using the spin column-based centrifugation, reversed transcribed and analysed for endogenous expressions of trib-1 and miR-202 using qPCR. One-ANOVA followed by Dunnett’s multiple comparison test was used to test for significance (P<0.05) across samples.

**Results:** It was observed that there was a significant decrease in trib-1 levels under these conditions of high glucose and cytokine stimulation and also with the combination of both whilst there was a consistent pattern of upregulation of MIR-202 under this conditions.

**Conclusion:** Taken together this study reveals that miR-202 is expressed in HepG2 cells, and a possible interaction between trib-1 and MIR-202 which could affect trib-1 stability and also the potentials for MIR-202 to be involved in some cellular activities in HepG2 cells relating to these conditions.

## 1. INTRODUCTION

The cardiovascular disease burden is a global one that affects both developed and developing countries. It is the leading cause of mortality in both men and women. Hyperlipidemia is classified as a major risk factor for the development of atherosclerosis a cardiovascular disease. An interplay between hyperlipidaemia and inflammation leads to the onset of atherogenesis. An increase in plasma levels of VLDL-C/LDL-C, as well as high circulating triglyceride, is considered atherogenic whereas the expression of HDL-C is “atheroprotective”^1^. Recent novel therapeutics for the treatment of cardiovascular disease in humans have been targeted towards the treatment of dyslipidaemia.

In the last couple of years, human genetic studies have identified several alleles that are robustly associated with coronary heart diseases, an increased risk factor for myocardial infarction and also other known complications of cardiovascular disease, among which includes polymorphisms located downstream of the human chromosome 8q24^2,3,4,5^. This gene locus contains the trib-1 gene. Following the results from genome-wide association studies, the trib-1 gene has been robustly linked with regulators of dyslipidemia such as; plasma triglyceride, low-density lipoprotein cholesterol, high-density lipoprotein cholesterol and cardiovascular risk^5,6,7^ as well as a regulator of lipid metabolism^8^. In a functional study, murine models of trib-1 knockout and liver-specific overexpression techniques established that the gene product of tribbles homolog 1 plays a mechanistic role in lipid metabolism^9^. Interestingly the trib-1 gene product has been reported to be highly unstable with a half-life of approximately one hour and as such expressed at low levels in body tissues where it is found^10^ and very little knowledge on how its expression is regulated. With evidence so far from previous studies, it is now apparent that the expression of trib-1 in the liver is “atheroprotective” and also essential for lipid metabolism which explains why it is associated with decreased risk for atherosclerosis as the liver is the major site for secretion and processing of circulating lipoproteins.

MicroRNAs are endogenous single-stranded non-coding RNAs of ∼22 nucleotides that affect gene expression at the post-transcriptional level by degradation of its messenger RNA or suppression of protein translation and are highly conserved in the human genome. Since their discovery about two decades ago in nematodes, more than a 1000 microRNAs have also been identified in the human genome and grossly distributed in different cells and tissues of the body, and are involved in various cellular activities and tissue differentiation^11^. Numerous microRNAs are associated with CVD & its risk factors, of these, a few of them like miR-130, miR-24, miR-122, miR 30c & miR-33 have been identified as important regulators of lipid metabolism^12,13,14,15,16,17^.

In a previous study, it’s been discovered that miR-202 can bind to the 3’ UTR of trib 1 gene which could cause mRNA degradation and reduce its expression^18^. In this present study, the expression levels of trib-1 gene and miR-202 in HEPG^2^ cells was examined under *in vitro* conditions that are considered to be similar to cases of hyperlipidaemia and high glucose as we understand very well that an interplay between hyperlipidaemia and inflammatory responses in the arterial wall is a causal risk factor for atherosclerosis.

## 2. MATERIALS AND METHODS

### Study Area

This study was carried out over a period of four (4) months.

### Cell Culture

HepG_2_ cells were obtained from the cardiovascular research unit, Royal Hallampshire Hospital, Sheffield and maintained in growth medium - Dulbecco’s Modified Eagle’s Medium-low glucose concentration-1g/L (DMEM, Invitrogen) and media supplemented with 10% FBS,100µg/ml of penicillin and streptomycin (10mls). HepG_2_ cells were incubated under standard culture conditions of 5% carbon dioxide at 37°C. Approximately, 4×10^5^ cells were then seeded into a 6-well plate and incubated for stimulation.

### Cell Stimulation

#### Cytokine Stimulation

Hyperlipidaemia is a condition that gives rise to a cascade of inflammatory responses^19^. To mimic this environment *in vitro* we stimulated hepG2 cells with interleukin-1β (100ng/ml). Cells were stimulated 24 hours after seeding and harvested after 6 hours for lysis and RNA extraction.

#### High Glucose Stimulation

Type 2 diabetes is linked with dyslipidaemia and also classified as one of the secondary causes of hyperlipidaemia^20,21^, patients with type 2 diabetes often end up with complications of CVD. A characteristic feature of type 2 diabetes is high levels of sugar (glucose) in the blood as a result of Insulin resistance^22^. To model this in a cellular environment, cells were exposed to 2mls of high glucose-DMEM (4.5g/L of glucose) supplemented with the foetal bovine solution and Penicillin/streptomycin for 48 hours.

#### Cell Lysis & RNA Isolation

This was carried out using 350µl of lysis buffer together with 3.5µl of β-mercaptoethanol for each well. The cell lysate was transferred into a collection tube for subsequent RNA isolation.

For analysis of endogenous expression of miR-202 and corresponding mRNA levels of trib-1, total and small non-coding RNA was isolated from the cells after stimulation using PureLink miRNA isolation kit according to manufacturer’s instructions.

#### Measurement of RNA concentration

A Nanodrop spectrophotometer was used to measure the RNA concentration of each sample together with its purity and integrity. This was quantified by absorbance at 260nm. RNA with good yield was qualified as RNA with an OD_260/280_ of >1.8.

### Reverse transcription (cDNA synthesis)

#### Total RNA

2µg of total RNA was reversed transcribed to the first-strand cDNA in a 25µl synthesis reaction using the Promega kit and M-MLV reverse transcriptase according to the manufacturer’s instruction.

#### MicroRNA

The cDNA synthesis for microRNA was performed using the TaqMan microRNA reverse transcription kit components according to the manufacturer’s instruction with a 100ng of microRNA. For detection of miR-202, a miR-202 specific stem-loop primer was used for cDNA synthesis. Reverse transcription was performed using Veriti thermal cycler with the thermal protocol set at 16°C for 30 minutes, followed by 42°C for 30 minutes and a final step of 85°C for 5 minutes and held at 4°C.

#### Quantitative Real-time PCR

Using the cDNA samples with the Taqman universal PCR master mix with FAM-dye-labelled probes and gene-specific primer sets for trib-1, miR-202(Stem-loop forward primer: ACACTCCAGCTGGGAGAGGTATAGGGCA and reverse primer: CTCAACTGGTGTCGTGGAGTCGGCAATTCAGTTGAGTTCCCATG for humans), qPCR analysis was conducted using CFX384 Real-Time System. Thermocycling was performed using the following conditions: 95°c for 10mins, 95°c for 10s, 60°c for the 30s for 40 cycles then 60°c for 60s. Relative gene expression was calculated using the comparative Ct (ΔΔCt) method and B-actin as a housekeeping gene. Values are presented as fold change relative to control cells.

## Statistical Analysis

All data are expressed as the mean ± standard error. Results were analysed by one-way analysis of variance; followed by Dunnett’s multiple comparison test using Graph pad prism 6.0 software (Graph pad Software, Inc.). P-values of <0.05 were considered to indicate a statistically significant difference.

## 3. RESULTS

### 3.1 miR-202 is expressed in hepG2 cells

The expression of miR-202 was analysed under metabolic and inflammatory conditions of treatment with high glucose for 48 hours and IL-1β stimulation for 6 hours and a combination of both treatments. The expression level of miR-202 was examined using real-time PCR with a miR-202 specific cDNA template. There was evidence of miR-202 expression in this cell line under these conditions, seen with an upregulation in fold difference of 1.2 – 2.0 between samples as compared to the control (Fig 1).

**Figure 1:**
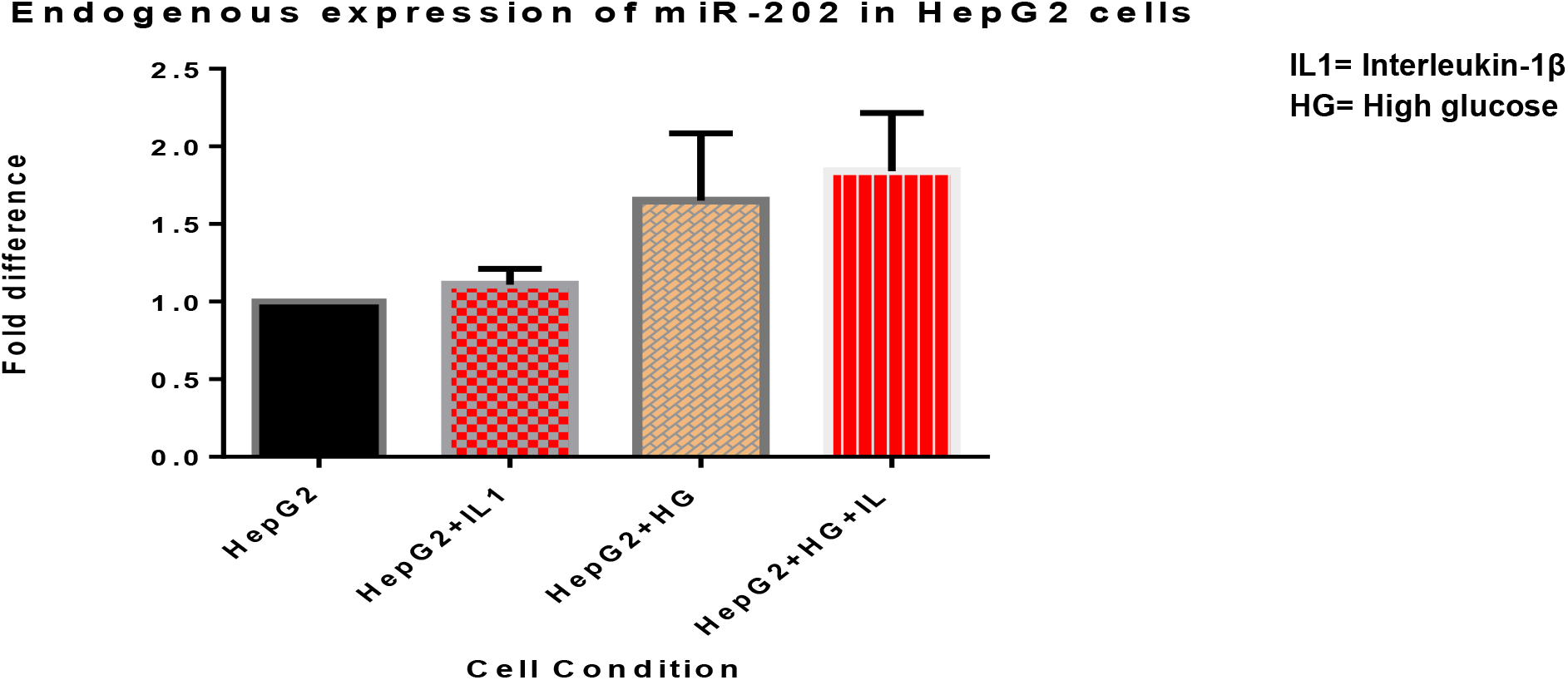
miR-202 expression in hepG2 cells. miR202 is expressed in hepG2 cells under conditions of metabolic and inflammatory stress. miRNA was isolated from hepG2 cells stimulated with IL-1β and cultured in 48 hours of high glucose and cDNA was synthesized using a miR-202 stem-loop specific primer. Expression levels of miR-202 were quantified using real-time PCR and analysed using one-way ANOVA followed by Dunnett’s multiple comparison test comparing all treated samples to the control.

### 3.2 Effects of endogenous miR-202 expression on the trib-1 level

To study the relationship between miR-202 and trib-1 under these cell type conditions, we also measured the mRNA levels of trib-1 in these cell samples stimulated with IL-1β and exposed to 48 hours of high glucose. Compared to the control, the trib-1 level was significantly reduced in the treated samples (P<0.05; Fig 2). Which was in contrast to the pattern observed of miR-202 expression.

**Figure 2:**
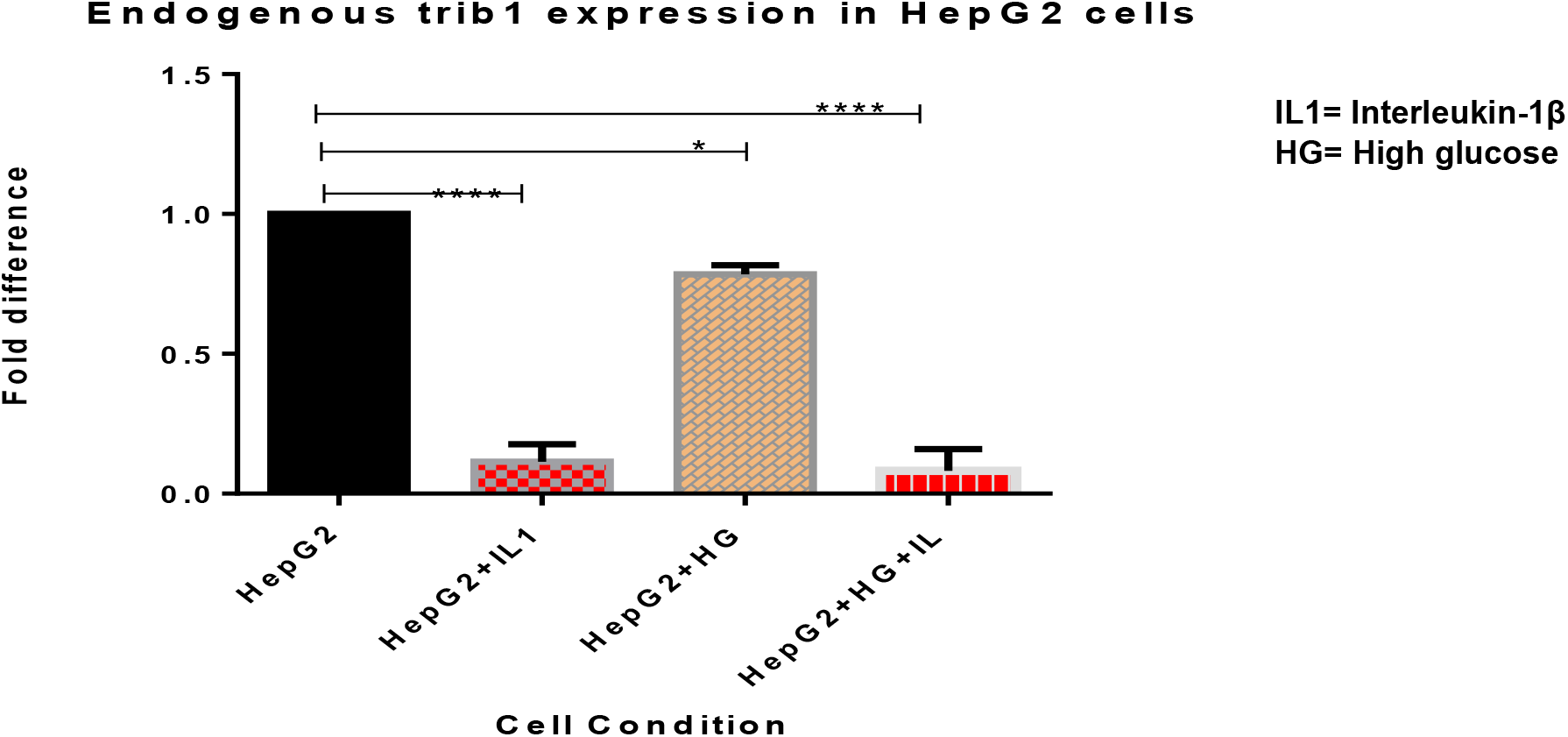
trib-1 expression decreases significantly under inflammatory and metabolic stress conditions. Total RNA isolated from HepG2 cells stimulated with IL-1β and cultured in 48 hours of high glucose was reversed transcribed and mRNA levels of trib-1 quantified using real-time PCR and analyzed using one-way ANOVA followed by Dunnett’s multiple comparison test (^****^P<0.05).

## 4. DISCUSSION

Data from this study reveals that mir-202 is expressed in hepG2 cells, its level of expression is also influenced, regulated, and affected by the in vitro cellular conditions of high glucose and cytokine stimulation. In this study, under the same cellular conditions, it is observed that trib-1 level is significantly down-regulated whereas there is an expression of mir-202. There were consistent changes in the pattern of trib-1 expression as it was with a mir-202 expression which was inversely proportional albeit under this cellular conditions of high glucose cytokine stimulation.

Genetic variations of trib1 locus have been linked to levels of plasma triglycerides, lipoproteins/cholesterol and risk of CVD^2,3,4,5,23,24,25,26,27,28^. Trib1 plays an important role in lipid metabolism as well as in human hepatocyte biology^6,8,29,30^. How trib-1 modulates hepatic lipogenesis is still in question, Ishizuka *et al*.,^31^ suggest that it could be through molecular interactions whereby it modulates genes involved in lipid metabolism, and recent studies reveal that the suppression of its expression dysregulates various fundamental pathways in liver functioning^6^, the exact mechanism for trib-1 function remains obscure.

The instability of trib-1 gene expression has been established with a half-life of less than an hour^10^ and equally very unstable in hepG2 cells^6,10^. Various genetic markers have been implicated in the regulation of gene expression, with the most recent being the discovery of small non-coding RNAs known as microRNAs. MicroRNAs are regulators of gene expression, they could function as master or key regulators of a process and could have different gene target which they can regulate all at the same time. A target gene could have more than one miRNA binding site and its expression could be regulated cooperatively by its various miRNA targets^32^. The roles of miRNAs in various human diseases have been characterized using microRNA profiling. Dysregulation of microRNAs is relevant in cancer, neurological, cardiovascular and other developmental diseases^32^.

miRNAs regulate gene expression in a sequence-specific manner by targeting 3’ UTR of mRNAs through partial or complete base pairing from their seed region. Bioinformatics search using miRNA target prediction algorithms had discovered a miR-202 binding site which overlaps in the trib-1 3’ UTR and this also correlates with studies conducted for target home profiling in lymphomagenesis, a RIP-Chip assay identifies trib-1 transcript as a potential gene target of miR-202^18^.

MiR-202 is located on chromosome 10q26^33^, its dysregulation has been highly associated with the pathogenesis of various human diseases especially cancer^18,33,34,35,36,37^.

In the light of all these findings, our study was aimed at understanding the mechanism of action of miR-202 in the regulation of hepatic trib-1. First, we evaluated miR-202 expression in a hepatic cell line (HepG2) since the liver is the major site of lipid metabolism. The *in vitro* conditions used were to enable us to analyse the expression of miR-202 under these conditions of metabolic and inflammatory stress, these conditions were selected under prevailing knowledge of the implication of hyperlipidaemia as inflammatory stress. It’s been reported that IL-1β induces hyperlipidaemia in rats and affects lipid metabolism enzyme leading to increased levels of serum lipid^5^ and high glucose which is symbolic of type2 diabetes. T2D has been identified as a secondary predisposing factor for hyperlipidaemia as insulin resistance causes dysregulation of plasma lipid/lipoprotein which are also described as metabolic defects in the body^38^. Evidently, from our observation, miR-202 was expressed in hepG2 cells (fig. 2) from the plotted bar graph in our results (fig. 2) miR-202 is expressed endogenously in hepG2 cells and, likely, it is also up-regulated under these conditions. Studies have reported low levels of expression of miR-202 in hepG2 (HCC cell lines)^33^, this makes it rather intriguing that we observed elevated patterns of miR-202 when compared to the control under these cellular conditions. It would be worthy to investigate this further, as this pattern of expression could probably be used as a biomarker for hepatic metabolic and inflammatory defects, as prolonged metabolic stress is linked with negative effects^21^, this could also be the case for miR-202 expression.

Looking at trib-1 expression levels under these same conditions, the total RNA used for this were also isolated from the same cell samples which we isolated the miRNA. We noticed a significant decrease in trib-1 mRNA levels as compared to the control (P<0.05). As much as it was not inversely proportional to miR-202 expression, it is almost similar to results obtained from *in vivo* studies were hyperlipidaemic conditions are associated with reduced levels of trib-1^2^ even though this study only quantifies trib-1 mRNA levels. Interestingly this could serve as an *in vitro* verification of existing data. Mouse trib-1 and human trib-1 have almost identical gene sequence, TargetScan release 6.2 algorithm based on evolutionary conservation shows that miR-202 binding sites on the trib-1 transcript are conserved in 20 other animal species including mouse species which has been used in previous studies of trib-1 function in lipid metabolism and could also be tested further in the light of this findings.

As much as it is said that Trib-1 levels expressed in HepG2 cells are comparable to those of primary hepatocytes, it has been established that trib-1 is unstable in HepG2 cells^10,39^. From our studies, we observed that the increasing changes in miR-202 level across these cellular conditions were associated with a significant reduction in trib-1 mRNA levels. Looking at the results (Figures 1 & 2) obtained from endogenous expression of miR-202 and trib-1, expression levels of miR-202 was upregulated across treatments while there was a significant reduction in trib-1 mRNA levels, and this correlates with miRNA studies where mRNA levels of target gene reduce as there is an overexpression of the potential miRNA^33,35,40^ and could be another confounding possibility for its low half-life in Hepg2 cells. Although, not reported an in vitro verification of this result was carried out using a luciferase plasmid construct. MicroRNA regulates gene expression by binding to 3’ UTR of its target mRNA and either cause degradation or suppresses translation. For this study we used luciferase plasmid construct with a truncated wild type trib-1 to explore this mode of action in HepG2 cells, our results showed that overexpressing miR-202 reduced trib-1levels by ∼50% and inhibiting miR-202 had reversed effects from the dual luciferase assay readings; although this result was not statistically significant. This effect also shows that miR-202 can bind to trib-1 in hepG2 cells.

## CONCLUSION

MiR-202 is a known target for trib-1 transcript, this study has provided evidence for its implication in atherosclerosis through its up regulated expression under conditions of inflammatory stress, metabolic stress and targeting of trib-1. Very little is known about the functions of trib-1 in lipid metabolism, however, glucose metabolism has been linked to trib-1. In this study increased glucose concentrations in hepg2 cells have also been seen to supress trib-1 level, this suppression has been linked to be consistent with its short half-life, we have also established that under this conditions, miR-202, a known culprit in post-transcriptional regulation of gene expression is also highly expressed.

Greater clarity of the mechanism of genetic regulation of trib-1 would greatly strengthen the debate for trib-1 based therapeutics.

## Significance Statement

This study has revealed that microRNA-202 a known target of trib-1 transcript is expressed in HepG2 an hepatic cell line under conditions that predisposes to hyperlipidaemia and inflammatory responses.

This study will other researchers to pay more attention to microRNAs in lipid metabolism and atherosclerosis.

## Acknowledgements

this study was based at the cardiovascular research unit of the department of infection, immunity and cardiovascular disease of the University of Sheffield medical school, we are grateful to them for providing the opportunity and support for this study.

